# Evolutionary genomics of peach and almond domestication

**DOI:** 10.1101/060160

**Authors:** Dianne Velasco, Josh Hough, Mallikarjuna Aradhya, Jeffrey Ross-Ibarra

## Abstract

The domesticated almond [Prunus dulcis (L.) Batsch] and peach [P. persica (Mill.) D. A. Webb] originate on opposite sides of Asia and were independently domesticated approximately 5000 years ago. While interfertile, they possess alternate mating systems and differ in a number of morphological and physiological traits. Here we evaluated patterns of genome-wide diversity in both almond and peach to better understand the impacts of mating system, adaptation, and domestication on the evolution of these taxa. Almond has ∼7X the genetic diversity of peach, and high genome-wide F_ST_ values support their status as separate species. We estimated a divergence time of approximately 8 Mya, coinciding with an active period of uplift in the northeast Tibetan Plateau and subsequent Asian climate change. We see no evidence of bottleneck during domestication of either species, but identify a number of regions showing signatures of selection during domestication and a significant overlap in candidate regions between peach and almond. While we expected gene expression in fruit to overlap with candidate selected regions, instead we find enrichment for loci highly differentiated between the species, consistent with recent fossil evidence suggesting fruit divergence long preceded domestication. Taken together this study tells us how closely related tree species evolve and are domesticated, the impact of these events on their genomes, and the utility of genomic information for long-lived species. Further exploration of this data will contribute to the genetic knowledge of these species and provide information regarding targets of selection for breeding application and further the understanding of evolution in these species.

## Introduction

*Prunus* is a large genus in the family Rosaceae with approximately two hundred species, including multiple domesticated crops such as almond, apricot, cherry, peach, and plum (Rehder, 1940; Potter, 2011). Peach [*P. persica* (Mill.) D. A. Webb] and almond [*P. dulcis* (L.) Batsch] are two of the three most economically important domesticates in *Prunus* globally, and share a number of similarities, including perenniality, precocity, and genome size and organization (Baird et al., 1994; Arús et al., 2012). However, the two species also have striking differences. While peaches are harvested for their indehiscent fleshy mesocarp, almonds are harvested for their seed, encased in a stony endocarp and a leathery, dehiscent mesocarp and exocarp (see Figure S1). And while almond, like most *Prunus* species, exhibits *S*-RNase based gametophytic self-incompatibility, peach is self-compatible (Hedrick et al., 1917; Wellington et al., 1929). Almond and peach also differ for other traits, such as life span (Gradziel, 2011), chilling requirements (Alonso et al., 2005; Dozier et al., 1990; Scorza and Okie, 1991), and adventitious root generation (Kester and Sartori, 1966).

Domestication of almond and peach occurred independently approximately 5000 BP in the Fertile Crescent and China (Zohary et al., 2012), respectively, followed by global dissemination beginning before 2300 BP (Hedrick et al., 1917; Edwards, 1975; Gradziel, 2011; Zheng et al., 2014). The few obvious domestication traits in almond are reduced toxicity, thinner endocarp, and increased seed size, while domestication in peach is characterized by diverse fruit morphology (size, color, texture, shape, etc.) and self-compatibility. Other traits not typically associated with domestication, such as precocity, adventitious rooting, graft compatibility, or tree architecture, may also have been selected during domestication or subsequent breeding (reviewed in Miller and Gross, 2011; Spiegel-Roy, 1986). Efforts to identify the wild progenitors of either almond or peach by examining species relationships within subgenus *Amygdalus* have produced inconsistent species trees and numerous polytomies (Mowrey et al., 1990; Browicz and Zohary, 1996; Ladizinsky, 1999; Aradhya et al., 2004; Bassi and Monet, 2008; Zeinalabedini et al., 2010; Verde et al., 2013). Given uncertainty in the wild progenitors and the difficulties associated with long generation times, QTL-mapping approaches to investigate peach or almond domestication are thus impractical. In contrast, comparatively fast and inexpensive sequencing makes population genetic approaches (cf. Ross-Ibarra et al., 2007) an attractive option, enabling the identification of domestication loci and study of the genome-wide impacts of changes in mating system.

Both domestication and mating system have been shown to shape genomic patterns of diversity in annual species (Glémin et al., 2006; Doebley et al., 2006; Hazzouri et al., 2013; Slotte et al., 2013), but the impacts of these forces on tree species remains poorly documented (McKey et al. 2010; Miller and Gross 2011; Gaut et al. 2015; but see Hamrick et al. 1992 for relevant analyses of allozyme diversity data). Mating system differences between closely related species pairs has been shown to significantly affect many aspects of genome evolution in *Arabidopsis*, *Capsella*, and *Collinsia*, including lower nucleotide diversity, higher linkage disequilibrium (LD), and reduced effective population size (N_*e*_) (Hazzouri et al., 2013; Slotte et al., 2013; Wright et al., 2013). Demographic bottlenecks associated with domestication may also reduce diversity genome-wide, and selection during domestication will reduce diversity even further at specific loci (Glémin et al., 2006; Doebley et al., 2006). While studies in perennials, particularly tree fruit crops, suggest they have lost little genetic diversity due to domestication (reviewed in Miller and Gross, 2011), recent analysis of resequenced peach genomes are consistent with lower genetic diversity and higher LD across the genome compared to related wild species (Verde et al., 2013; Cao et al., 2014). No such genome-wide analysis of diversity in almonds currently exists, however, and little is known about how differences in mating system affect changes in diversity during domestication.

Here we leverage both new and published genome sequences to present an evolutionary genomic analysis of the effects of domestication and mating system on diversity in both almond and peach. Understanding the impact of mating system will expand the basic knowledge of genome evolution in a perennial species pair with contrasting mating systems, and identification of candidate domestication loci will provide an opportunity to assess convergence during domestication and compare tree domestication to that of annual crops.

## Materials and Methods

### Samples

We used 13 almond and 13 peach genomes for all analyses, which included both public and newly resequenced data (Tables 1, S1). In addition, we used one peach-almond F_1_ hybrid and one peach with Nonpareil almond in its pedigree as checks for admixture analysis. For this study we resequenced nine almonds, one peach, an almond-peach F_1_ hybrid, and the plum *P. cerasifera* as an outgroup (Tables 1, S1). Fresh leaves and dormant cuttings collected from multiple sources were either desiccated with silica or stored at 4C prior to DNA isolation. We isolated DNA following a modified CTAB method (Doyle, 1987).

**Table 1:** *P. dulcis*, *P. persica* and outgroup species used in analyses.

| Species | *n* | Avg. Depth | Reference |
| --- | --- | --- | --- |
| *P. dulcis* | 4 | 7.76 | Koepke et al., 2013 |
| *P. dulcis* | 9 | 19.34 | this study |
| *P. persica* | 10 | 19.13 | Verde et al., 2013 |
| *P. persica* | 2 | 13.78 | Ahmad et al., 2011 |
| *P. persica* | 1 | 37.36 | this study |
| *P. cerasifera* | 1 | 35.02 | this study |

Libraries for eight of the almond samples were prepared at UC Davis. We quantified the sample DNA with Quanti-iT Picogreen dsDNA assay (Invitrogen, Life Technologies) and then fragmented 1 *µ*g with a Bioruptor (Diagenode) for 11 cycles of 30 seconds ON and 30 seconds OFF per cycle. The resulting DNA fragment ends were then repaired with NEBNext End Repair (New England BioLabs) followed by the addition of deoxyadenosine triphosphate to the 3-prime ends with Klenow Fragment (New England BioLabs). We then ligated barcoded Illumina TrueSeq adapters (Affymetrix) to the A-tailed fragments with Quick Ligase (New England BioLabs). Between each enzymatic step we washed the DNA with Sera-Mag SpeedBeads (GE Life Sciences, Pittsburgh). The resulting libraries were quantified with a Qubit (Life Technologies) and sized using a BioAnalyzer DNA 12000 chip (Agilent Technologies). Libraries were sent to UC Berkeley (Berkeley, Qb3) for quantification by qPCR, multiplexing, and sequencing for 100 bp paired-end reads in a single HiSeq 2000 (Illumina) lane. DNA from the remaining samples (Tables 1, S1) was submitted to BGI (Shenzen, China) for library preparation and sequenced using 100 bp paired-end reads at their facility in Hong Kong. Sequence data are available from SRA (http://www.ncbi.nlm.nih.gov/sra) and the associated run numbers are given in Table S1.

### Analysis

#### Quality Control and Mapping

All FASTQ files were trimmed of remnant adapter sequences using Scythe (github.com/vsbuffalo/scythe) and then further trimmed for base quality with Sickle (github.com/najoshi/sickle) using default parameters for both. Trimmed reads were then aligned to the *P. persica* v1.0 reference (www.rosaceae.org) using BWA-MEM (Li, 2013) with a minimum seed length of 10 and internal seed length of 2.85. After filtering for a minimum mapping quality of 30 and base quality of 20, sequence depth averaged 15.8X (4.7X to 34.6X) in almond and 19.7X (11.2X to 35.4X in peach; Table S1, Figure S2).

#### Diversity and Candidate Gene Identification

We estimated inbreeding coefficients using *ngsF* in the *ngsTools* suite (Fumagalli et al., 2014), and then calculated genotype likelihoods in ANGSD (Korneliussen et al., 2014) incorporating our inbreeding estimates. We calculated several population genetics statistics, including pairwise nucleotide diversity (*θ_π_*; Nei and Li, 1979), Tajima’s *D* (*D*; Tajima, 1989), Fay and Wu’s *H* (*H*; Fay and Wu, 2000), and Zeng’s *E* (*E*; Zeng et al., 2006) using the *thetaStat* subprogram in ANGSD. Diversity values were estimated in overlapping 1000 bp windows with 50 bp steps, removing windows with less than 150 bp of sequence after filtering. Additionally we calculated a normalized *θ_π_* value by dividing per window *θ_π_* by mean *θ_π_* in each species. To identify candidate genes possibly selected during domestication, we filtered for genes in the lowest 5% empirical quantile of each diversity statistic. We further analyzed candidate loci for gene ontology (GO) using *P. persica* protein gene identifiers in the singular enrichment analysis tool and Fisher’s exact test using default statistical options at the AgriGO website (http://bioinfo.cau.edu.cn/agriGO/).

#### Population Comparisons

We treated peach samples and almond samples as two populations to evaluate population structure. We performed a principal component analysis (PCA) with *ngsPopGen* (Fumagalli et al., 2014) and used *NGSadmix* (Skotte et al., 2013) to perform an admixture analysis and assign proportions of almond and peach population to individuals using *K* = 2 through *K* = 6 as the number of potential subpopulations. Finally, we examined population differentiation by estimating *F*_*ST*_ genome-wide and in sliding windows (1000 bp windows with a 50 bp step) after removing windows with *<* 150bp of sequence.

#### Estimating historical changes in N_*e*_

To model the history of these species and infer the historical changes in effective population size that may have occurred prior to or during domestication, we analyzed peach and almond samples using the Multiple Sequentially Markovian Coalescent (MSMC) method (Schiffels and Durbin, 2014). This approach uses the observed pattern of mutations in multiple individuals to infer the time to the most recent common ancestor between pairs of sampled alleles, and provides maximum-likelihood estimation of population size as a function of time. Using the msmc software (github.com/stschiff/msmc) and msmc-tools (github.com/stschiff/msmc-tools), we applied this method to 10 individuals from our study (five peach and five almond samples; peach: PP02, PP03, PP04, PP05, PP13; almond: PD03, PD04, PD05, PD06, PD07) in two separate analyses. For each individual, we first identified SNPs for each chromosome using samtools mpileup (v. 1.3.1) with a minimum mapping and base quality cut off of 20. We filtered sites for depth ¡15 using VCFtools (v. 0.1.13), and removed indels using bcftools (v. 1.3.1). To estimate population size changes during the recent past (since domestication), we ran the full MSMC model for peach and almond separately using the combined set of five samples for each run. To estimate changes in *N*_*e*_ over a longer time period (2 mya), we applied the PSMC’ model (see Schiffels and Durbin, 2014) to each sample individually. To convert the mutation-scaled coalescent times and population sizes obtained from these analyses, we divided by a mutation rate of *µ* = 10^*−8*^ mutations per nucleotide per generation, and assumed a generation time of 10 years for both peach and almond. The models and inference algorithms for PSMC’ and MSMC are available from github.com/stschiff/msmc, and our code for analyzing peach and almond samples is available from https://github.com/houghjosh/peach.

#### Gene Expression

We downloaded ten SRA RNA-seq runs from four peach and almond tissues (Table S2). All runs were from either general transcriptome sequencing (Jo et al., 2015) or controls of differential expression experiments (Wang et al., 2013; Mousavi et al., 2014; Sanhueza et al., 2015). We then converted the runs into their paired FASTQ files using SRA-toolkit (v. 2.3.4) and quantified expression for each run separately against the peach transcriptome (v. 1.0) using kallisto (Bray et al., 2016). For each sequencing run kallisto outputs the transcripts per million (TPM), a within library proportional measurement, for each gene. Each gene was then annotated with its candidate or non-candidate status based on F_*ST*_, *θ_π_*,

Tajima’s *D*, Zeng’s *E*, or Fay and Wu’s *H* for both almond and peach. We also calculated the number of tissues in which each gene was expressed and the mean expression level in each tissue (across runs in which the gene was expressed).

## Results and Discussion

### Diversity

Genome-wide nucleotide diversity (*θ_π_*; Figures S5 and S6) in almond is nearly sevenfold higher than in peach (0.0186 and 0.0027, respectively), and these differences were more pronounced in non-genic regions of the genome (Tables 2 and S4). Though differences in diversity between peach and almond have been known from analyses using multiple marker systems (Mowrey et al., 1990; Byrne, 1990; Martínez-Goómez et al., 2003; Aradhya et al., 2004), this study is the first comparison of whole genome sequences using multiple diverse individuals from both species. Previous genome scans of peach found low levels of genetic diversity compared to the closely related wild species, *P. kansuensis*, *P. mira*, and 16. *davidiana* (Verde et al., 2013; Cao et al., 2014). Of these, only *P. davidiana* is outcrossing, and Verde et al. (2013) found it had the greatest nucleotide diversity of the species they examined, approximately three-fold higher than domesticated peach. Despite its domesticated status, almond retains more genetic diversity than any of the peach species studied thus far, suggesting that mating system explains more of the differences in diversity among species than domestication. Finally, we observed considerable variation in diversity statistics among chromosomes in both species, including up to two-fold differences in nucleotide diversity in peach (Table S4), perhaps suggesting the relatively recent effects of selection during domestication.

**Table 2:**
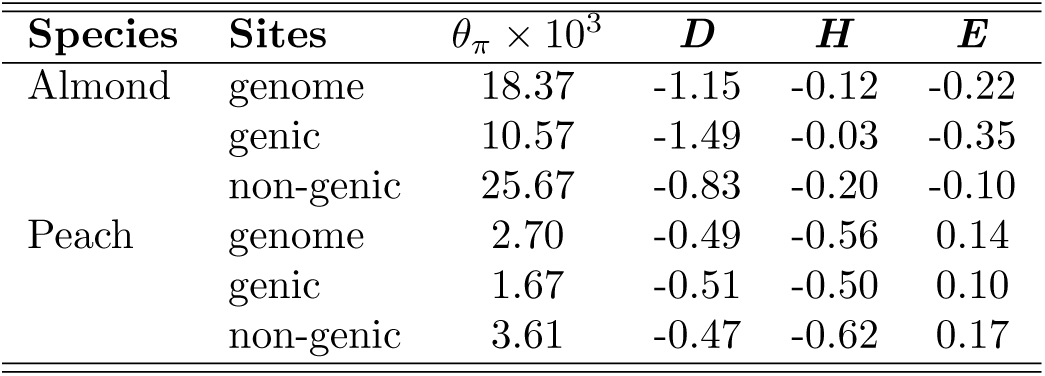
Genome-wide, genic, and non-genic diversity statistics and neutrality test values.

Mean values of Tajima’s *D* were negative for both almond and peach (Table 2), suggesting a genome-wide excess of rare variants likely consistent with a history of population expansion. Strongly negative values of Tajima’s D have recently been reported in *Populus tremuloides*, a species also inferred to have undergone post-glacial population expansion in the Quaternary Wang et al. (2016). While the wild progenitors of almond and peach are not definitively known, the current range of wild almond species is much larger than that of wild peach taxa, perhaps reflecting either contrasting initial population sizes or differential expansion of ancestral progenitors during interglacial periods following the Last Glacial Maximum (20 KBP; LGM).

#### Historical changes in N_*e*_

To investigate the demographic factors that may have contributed to the strong allele frequency skews that we observed in both peach and almond (Table 2), we conducted a whole-genome analysis of coalescent rates between haplotypes through time using MSMC (Schiffels and Durbin, 2014). The results from this analysis provide the first detailed comparisons of demography in both peach and almond, and enabled us to obtain estimates of population size changes from approximately 2 million years ago up to ≈ 1000 years ago (i.e., the last 100 generations; Figure S8). We found no evidence for a domestication-associated population bottleneck in either peach or almond S8A. Instead, our results suggest that almond experienced a population expansion following a bottleneck ≈ 20,000 years ago, consistent with our observations of a strongly negative Tajima’s D and perhaps due to rapid human-mediated dispersal from east Asia (Delplancke et al., 2012). In peach, our results suggest a gradual decline in *N*_*e*_ beginning ≈ 2 mya (Figure S8B), and extending to 5000 years ago, after which the effective population size remains very low. Although our results do not support a bottleneck in peach, the gradual decline in *N*_*e*_ starting in the distant past (≈ 2 mya; Figure S8B) is consistent with the low overall diversity we observe (Table 2), and may reflect a shift to a higher selfing rate (Charlesworth, 2003).

Overall, our analyses suggest that although population bottlenecks or extreme population expansions have occurred during domestication in many crop species (Meyer et al., 2012; Beissinger et al., 2016), neither peach nor almond appear to have experienced such events. In this respect, our results mirror those from other domesticated woody perennial crop species, including grape and apple, which are also reported to lack domestication bottlenecks but maintain much of their ancestral genetic diversity (Myles et al., 2011; Gross et al., 2014). This difference between annual and perennial domesticated crops may be due to the unique life cycle features of perennials, including a long generation time, overlapping generations, a typically outcrossing mating system, as well as a more recent period of domestication (Gaut et al., 2015). That we also found a large reduction in *N*_*e*_ and neutral diversity in peach despite no evidence for a population bottleneck also highlights the possibility that, within woody perennials, mating system differences may play an important role in determining the propensity of these species to domestication-associated bottlenecks.

#### Inbreeding

We estimated the average inbreeding coefficient (*F*) for almond and peach to be 0.002 (0.000 to 0.027) and 0.197 (0.000 to 0.737), respectively (Table S3). Although two self-compatible almond varieties are included in this study, none of our almond samples are derived from self-fertilization, supporting the low estimated inbreeding values. Peaches in general are self-compatible (with the exception of male-sterile varieties), and three of the peach varieties sampled (PP06, PP08, and PP15) have inbreeding values consistent with self pollination in the preceding generation (*F* =0.74, 0.53, and 0.56, respectively). Consistent with its known history as the result of open-pollination (Hedrick et al., 1917), the Georgia Belle peach variety sampled was estimated to have *F* = 0.

While the estimated inbreeding value for almond is not unexpected given that it is self-incompatible, the average for peach is lower than previously estimated selfing rates (*s*) of 0.5 *−* 0.86 (*F* = 0.33 *−* 0.75 from 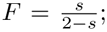 Fogle and Dermen 1969; Fogle 1977; Miller et al. 1989; Akagi et al. 2016). While the widely cited Miller et al. (1989) estimate was based on a single isozyme marker and is thus unable to separate self-fertilization with outcrossing to close relatives, the Akagi et al. (2016) estimate based on 5180 SNP markers is also high (*s* = 0.50 − 0.68 from *F* = 0.33 − 0.52). Our estimates are much closer to those from Aranzana et al. (2002), who estimated *s* = 0.148 (*F* = 0.08) from 35 microsatellites. In addition to differences in marker systems, these discrepancies are likely due at least in part to sampling, with estimates from outcrossed pedigrees (Aranzana et al., 2002) lower than those from landraces (Akagi et al., 2016). Broad examination of pedigree records, however, suggests our estimate of inbreeding is likely reasonable, as more than 67% of the 600 peaches in Okie (1998) were the result of outcrossing (Aranzana et al., 2002), including several of the varieties sampled here (Hedrick et al., 1917).

#### Population structure

Genome-wide, our data are consistent with previous estimates (Aradhya et al., 2004) in finding strong genetic differentiation between almond and peach (weighted *F*_*ST*_ = 0.605, Table S4, Figure S7). Like *F*_*ST*_, PCA also clearly distinguished almond from peach samples, primarily along PC1 (Figure 1). However, while PC2 and PC3 provided no further separation of peach samples they do allow further separation of almond samples (Figure 1).

**Figure 1:**
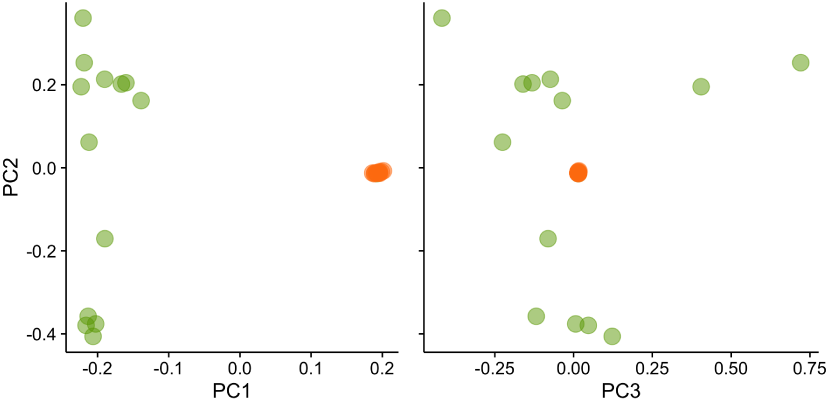
Principle component analysis of almond (green) and peach (orange).

Admixture analysis clearly assigns individuals to either almond or peach populations at *K* =2 (green and orange, respectively), including the correct identification of PD01 as an almond-peach F1 hybrid (Figure 2). Peach sample PP12, in contrast, should show approximately 12.5% almond based on its pedigree (Fresnedo-Ramírez et al., 2013) but in this analysis does not differ from other New World peaches in its assigned proportions. The fact that PP12 shows fewer total variants than PP13 (’Georgia Belle’; Fresnedo-Ramírez et al. 2013) is also inconsistent with recent almond ancestry, suggesting possible errors in the recorded pedigree.

**Figure 2:**
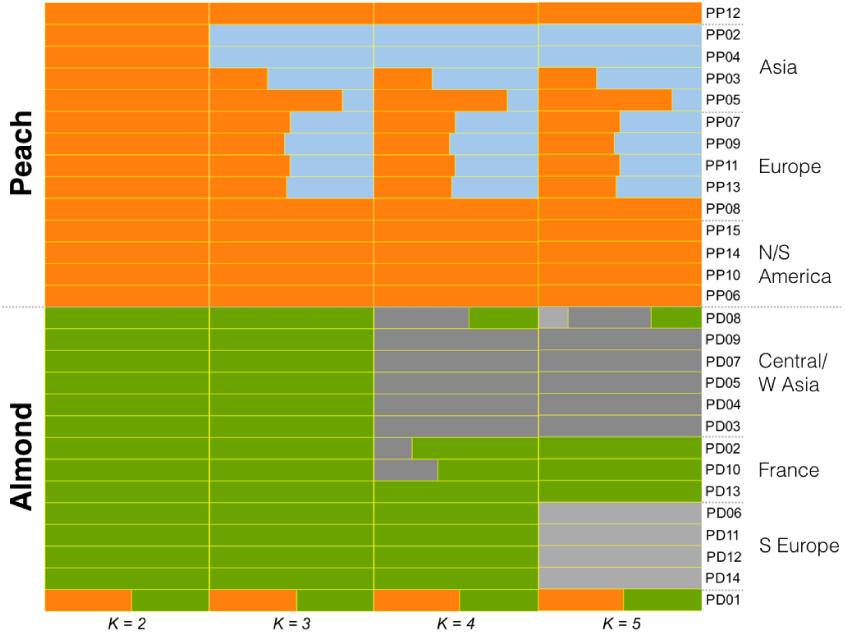
Admixture proportion of almond (PD) and peach (PP) for *K* =2 through *K* =5. With the exception of the purported hybrids, PD01 and PP12, sample origins generally correspond with an east (top) to west (bottom) orientation for each type (Table S1)

Increasing the number of clusters (*K*), we find evidence for population substructure in both peach and almond (Figures 2,S4) distinguished by geographic origin or breeding status. In the admixture plot (Figure 2), within both almond and peach groups, samples at the top have more eastern origins (Central Asia or China, respectively), whereas those towards the bottom have more western origins (Spain or New World, respectively). Almond samples from China, Pakistan, Iran, and Turkey (PD09, PD07, PD05, PD04 and PD03) group together at both *K* =4 and *K* =5. At *K* =5 a Mediterranean group of Italian and Spanish samples (PD06, PD11, PD12, and PD14) is identified, perhaps reflecting gene flow from North Africa into Spain and Italy (Delplancke et al., 2013). At *K* =6 PD01 forms a unique cluster and several other almonds shift assignments, suggesting an overestimation of the number of subgroups (Figure S4). Similar overall patterns of structure in peach and almond were found in previous studies (Li et al., 2013; Micheletti et al., 2015; Shen et al., 2015; Delplancke et al., 2013) as well, suggesting the use of local varieties as founders, limited exchange between Asian and European breeding programs, or recent utilization of diverse genetic resources is not reflected in the sampling. The foundations of most modern almond breeding programs began within the past century, due in part to the challenges of understanding self-incompatibility, whereas the self-compatible peach has had more widespread efforts directed towards its development for millenia (though western breeding increased or intensified only within the past 10 to 20 generations).

All of our analyses of differentiation provide unequivocal evidence distinguishing almonds from peaches, strongly supporting their status as distinct species. Previous molecular analyses have estimated a broad range of divergence times between these species, from 2.5 Mya (Vieira et al., 2008) to more than 47 Mya (Chin et al., 2014). One compelling idea for the origin of peach and almond is that climatic changes after Himalayan orogeny and Tibetan Plateau uplift led to isolation of an hypothesized ancestral species resulting in allopatric divergence of peach from almond (Chin et al., 2014). Consistent with this possibility, our estimates of *F*_*ST*_ and nucleotide diversity give a divergence time of ≈ 8 million years under a simple model of divergence in isolation (cf Holsinger and Weir, 2009) and assuming a mutation rate of *µ* = 10^*−8*^ per nucleotide and generation time of ≈ 10 years. This corresponds to a period of climatic change following significant geologic activity and uplift specifically in the northeastern section of the Tibetan Plateau (Fang et al., 2007; Molnar et al., 2010).

#### Candidate Loci

We next scanned the genomes of both almond and peach for potential candidate genes targeted by selection during domestication. In the lowest 5% quantile of Zeng’s E, we found 1334 and 1315 genes in peach and almond, respectively. Of these, peach and almond share 104, nearly double that expected by chance (permutation p-value *<* 0.001) and suggesting convergence in the process of domestication. In almond, candidate genes showed enrichment for gene ontology (GO) categories related to protein amino acid phosphorylation, ATP biosynthetic processes, regulation of ADP ribosylation factor (ARF) protein signal transduction, membrane and nucleus cellular components, ATP binding, ATPase and protein tyrosine kinase activities, and zinc ion binding; candidate genes in peach showed enrichment for the GO category related to cellular catabolic processes. We also identified the 1314 genes showing the greatest differentiation between species (top 5% quantile of *F*_*ST*_) but while these genes were enriched for a number of GO categories (Table S5) no clear patterns emerged.

We first investigated the *S*-locus in order to examine a genomic region known to differ between almond and peach both in sequence and function (Tao et al., 2007; Hanada et al., 2014). The *S*-locus controls gametophytic self-incompatibility in *Prunus* (reviewed in Wu et al. 2013). The *S*-locus haplotype block consists of two genes, *S*-RNase and the *S*-haplotype-specific F-box (*SFB*), which function in the pistil and pollen, respectively. In our data, the intergenic region 3’ to both the *S*-RNase and *SFB* loci shows elevated differentiation with one extremely high peak and low nucleotide diversity in peach (Figure 3A), observations consistent with recent work showing peach having only five known *S*-haplotypes, two of which have identical *SFB* alleles (Tao et al., 2007; Hanada et al., 2014).

**Figure 3:**
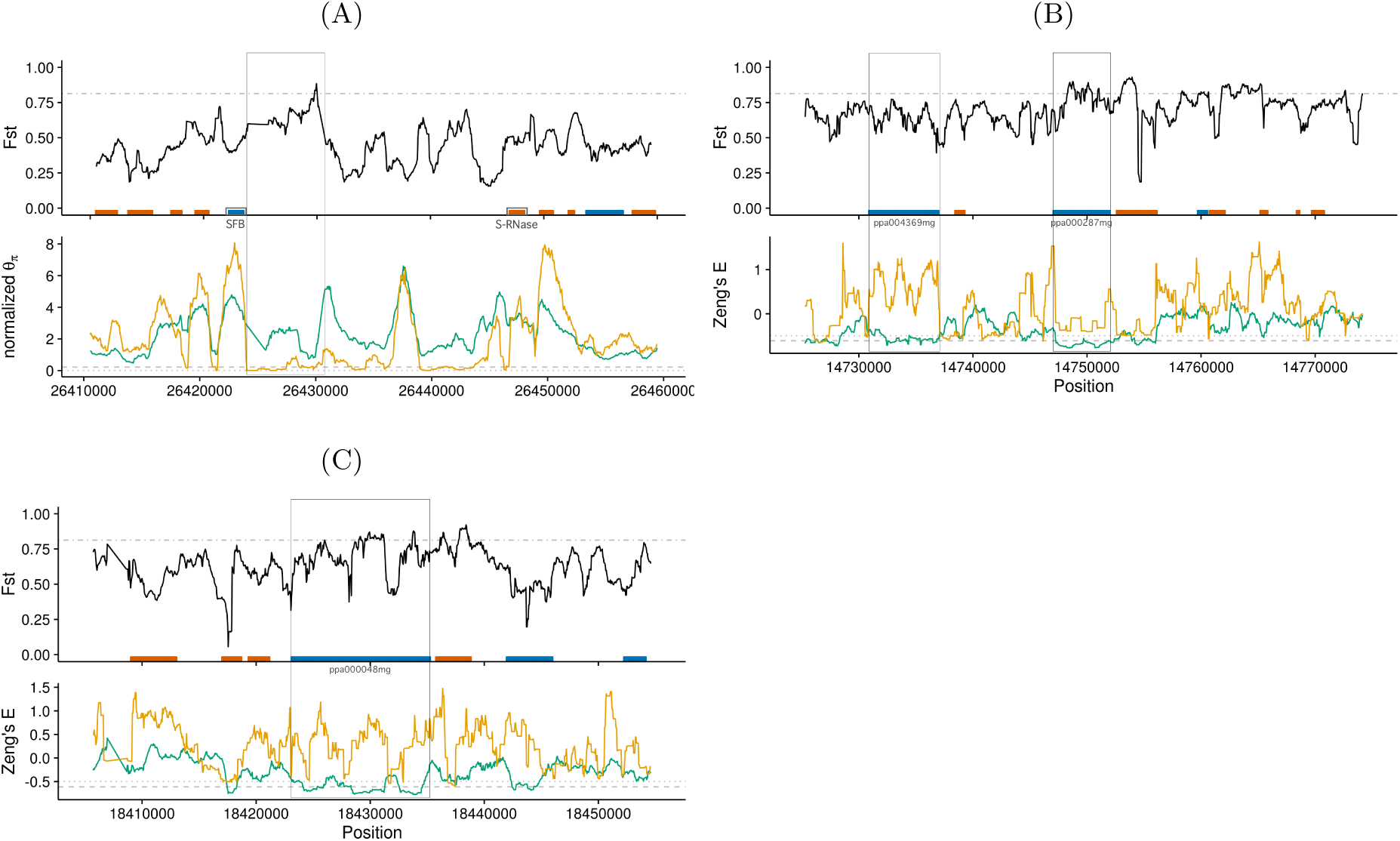
Select 50 Kb windows of the genome with high divergence (*F*_*ST*_) and either low normalized *θ_π_*(A) or Zeng’s E (B,C) of almond (green) and peach (orange). Genes annotated in the peach reference genome are represented in the *F*_*ST*_ plot by boxes colored by their location on the plus strand (blue) or minus strand (red). In the *F*_*ST*_ plots, the grey lines indicate the upper 5% threshold, whereas in the *θ_π_* and Zeng’s E panels the grey lines indicated the lower 5% thresholds of almond (dashed) and peach (dotted). Regions of interest, as described in the text, are boxed across adjacent panels and genes labeled. **A**. *S*-locus divergence and diversity with *S*-locus genes, *SFB* (blue) and *S*-RNase (red), located on opposite sides of the central gap. Diversity in peach is drastically reduced immediately 3’ to *SFB* but only somewhat reduced 3’ to *S*-RNase as might be expected for a linked locus. **B** & **C**. Loci of interest on chromosome 3.

Windows in the lowest 5% quantile of the summary statistics investigated were generally enriched for genic regions of the genome in both taxa, but the signal in peach was weak and enrichment was not consistent across all statistics evaluated (Table S6). Nonetheless, a number of individual regions genome-wide showed strong signatures of selection. We examined 50 Kb regions with contiguous windows in the bottom 5% quantile to focus our investigations of candidate genes. We focused on regions in both species for which there were overlapping regions of high *F*_*ST*_ and low *θ_π_* or Zeng’s *E* as these were significant for both peach and almond (permutation p-values 0 0.034; Table 3).

**Table 3:** Permutation probability for the overlap of neutrality test or *θ_π_* selected candidate genes with high *F*_*ST*_ selected candidate genes.

| Species | Tajima's $D$ | Fay & Wu's $H$ | Zeng's $E$ | $\theta_\pi$ |
| --- | --- | --- | --- | --- |
| Almond | 0 | 0 | 0 | 0 |
| Peach | 0.5854 | 0.3336 | 0.0342 | 0 |

While many intergenic and putative regulatory regions also showed interesting patterns in diversity statistics, we examined two regions of chromosome 3 with moderate to high *F*_*ST*_ and divergent values of Zeng’s E between peach and almond, specifically low values of Zeng’s E in almond (Figures 3B, 3C). The first of these regions (Figure 3B), contains the uncharacterized genes ppa004369mg (position 3:14730867.14736998; Uniprot identifier M5WRK6 PRUPE) and ppa00287mg (position 3:14747030.14752018; Uniprot identifier M5WX95 PRUPE), which have similarity to *γ*-aminobutyrate (GABA) transaminases in *Malus domesticus* and Myosin-1 in *Gossypium arboreum*, respectively. GABA is involved in signaling and nuclear regulation of cell wall modification and cell death through repression and activation, respectively, while GABA transaminases degrade GABA in the mitochondria and are reported to have a role in pollen-pistil interactions. Myosins are cellular motor proteins that act in concert with actin filaments for intracellular transport and cellular structure. The second region of interest on chromosome 3 (Figure 3C), contains the uncharacterized gene ppa000048mg (position 3:18423078.18435264, Uniprot identifier M5XGZ7 PRUPE). This gene is in the GO category of protein N-linked glycosylation and though it has high protein BLAST similarity among many species, few were annotated. Further investigation of additional regions with limited homology to characterized genes or functional information may be warranted given the poor characterization of genes in these species.

Given the importance of fruit morphology in peach we hypothesized that selection during domestication and subsequent breeding may have targeted genes primarily expressed in fruit tissue. To test this hypothesis, we compared gene expression in four tissues (peach fruit and leaf and almond ovary and anther) to candidate gene status. Candidates were over-represented among genes expressed in all tissues, and we saw no evidence of enrichment for tissue-specific expression in any of the four tissues (*χ*^2^ test showed significant under-enrichment in most cases; Table S7). Even among genes showing tissue-specific expression, we found no difference in expression between domestication candidates and non-candidates. We did, however, find that genes showing strong differentiation between almond and peach (highest 5% tail of *F*_*ST*_) showed higher levels of expression in both leaves and fruit. While we have no clear *a priori* hypothesis predicting differences in leaf-specific expression, higher fruit-specific expression among *F*_*ST*_ is certainly of note given the striking differences in fruit morphology between the species.

Contrary to our predictions, we find no evidence that domestication candidates are enriched for genes showing unusual patterns or levels of expression. Recent results, however, suggest that larger fruits may have much predated domestication. Seeds of a 2.6 My fossil peach *P. kunmingensis* were recently reported to be nearly identical to modern peaches (Su et al., 2015), and the observed correlation between seed size and fruit size in these taxa (Zheng et al., 2014) suggest fruit size was likely larger as well. Our finding that fruit-specific genes showing the strongest differentiation between species are more highly expressed is thus at least consistent with the possibility of selection for differences in fruit morphology between peach and almond predating domestication.

## Conclusions

One of the primary questions regarding domestication of perennial crops, particularly tree crops, is its genetic basis (Miller and Gross, 2011). Here we have examined two closely related domesticated tree species with alternate mating systems in an attempt to tease apart the genomic signatures of domestication and mating system and better understand these processes in perennial species. In addition to presenting evidence consistent with mating system effects in determining overall patterns of genetic diversity, our results identify numerous genes and genomic regions showing evidence of selection, provide evidence of convergence in the domestication of almond and peach, and fruit was not preferentially targeted during domestication but likely selected much earlier during species divergence. Finally, the high-coverage sequence we provide for a number of important cultivars may be useful to breeders and geneticists in identifying the causal basis of quantitative trait loci or developing marker sets for marker-assisted selection or genomic prediction.

## Acknowledgments

We thank Anne Lorant for almond sequencing library preparations and Emily Josephs, Michelle Stitzer, and two anonymous reviewers for their helpful comments and suggestions on the manuscript. Support for DV provided by the McDonald Endowment for UC Davis Plant Sciences Graduate Student Research Assistantship and the Almond Board of California (ABC; grant HORT16-Aradhya/Ledbetter). Resequencing funded by the ABC (grant HORT16-Aradhya/ Ledbetter) and a Henry A. Jastro Research Fellowship. Resequencing funded by the Henry A. Jastro Research Fellowship used the Vincent J. Coates Genomics Sequencing Laboratory at UC Berkeley, supported by NIH S10 Instrumentation Grants S10RR029668 and S10RR027303. J.R.I and J.H. were supported by funding from NSF Plant Genome project IOS-1238014.

---

**Figure S1:**
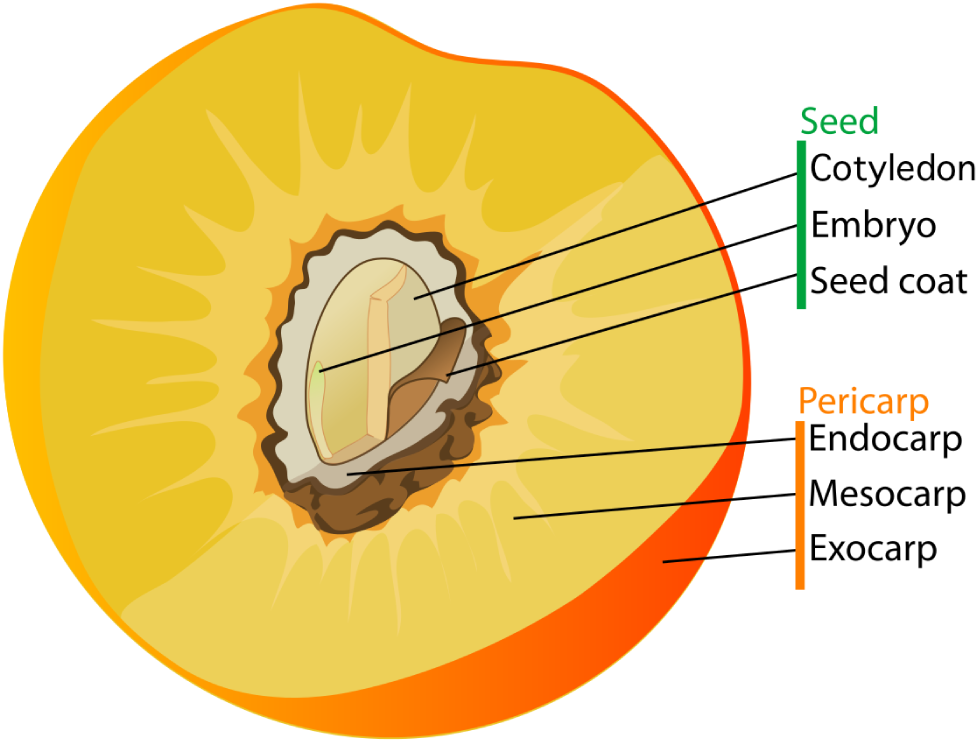
Peach and almond fruit and seed anatomy.

**Figure S2:**
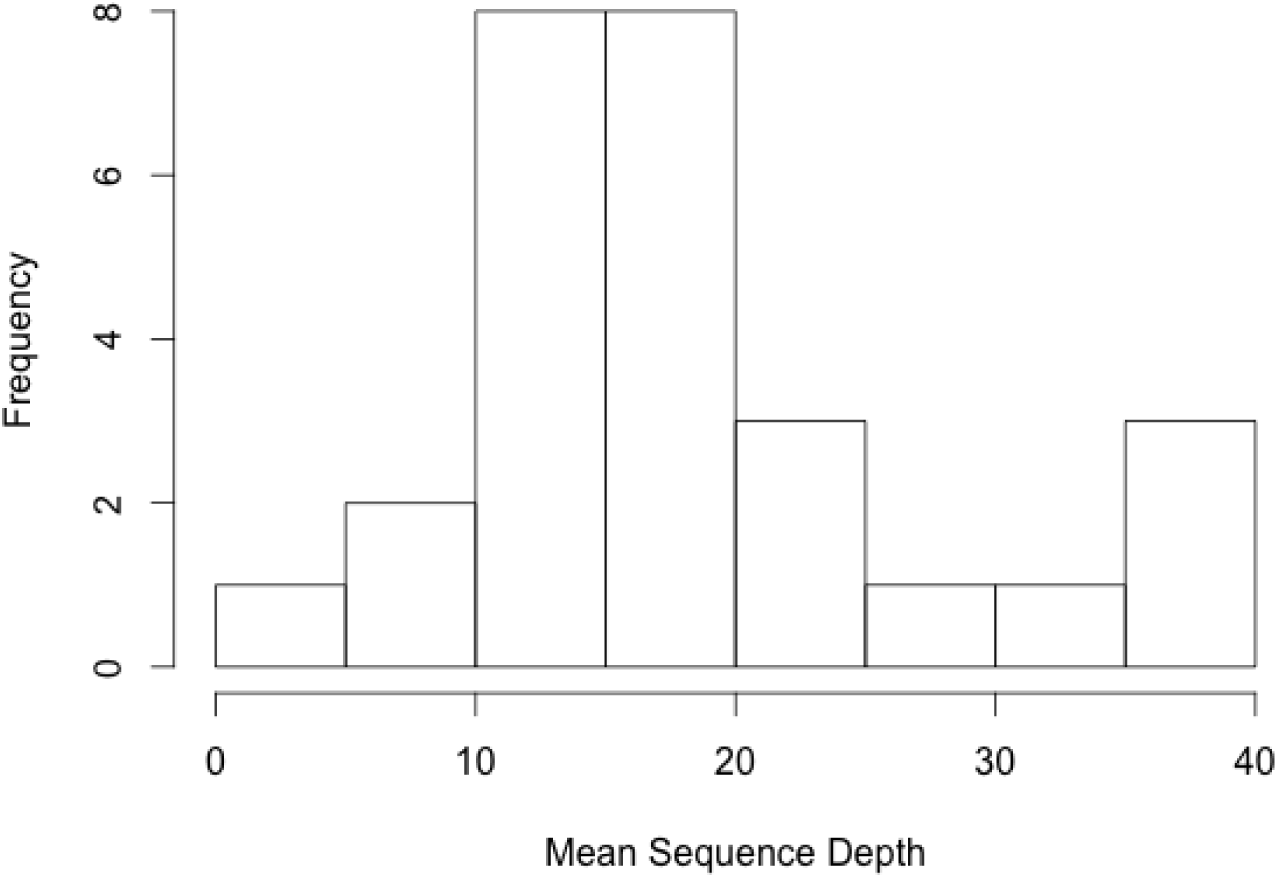
Mean mapped depth of peach and almond sequences used in this analysis filtered for mapping quality (MAPQ) scores *≥* 30 and base quality scores *≥* 20.

**Figure S3:**
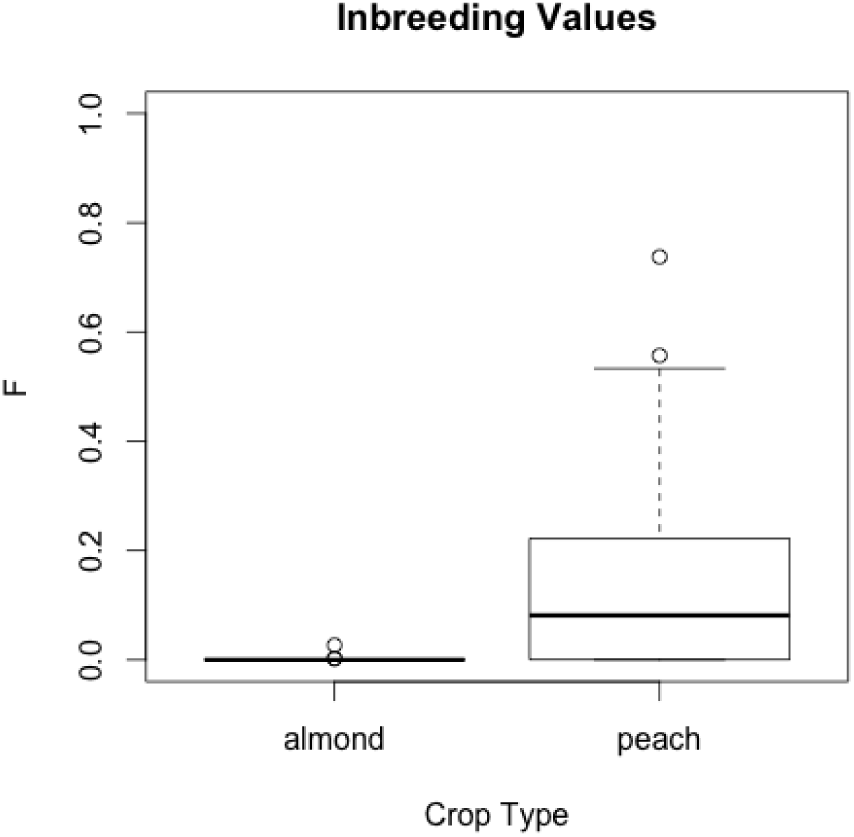
Distribution of inbreeding values for almond and peach samples studied.

**Figure S4:**
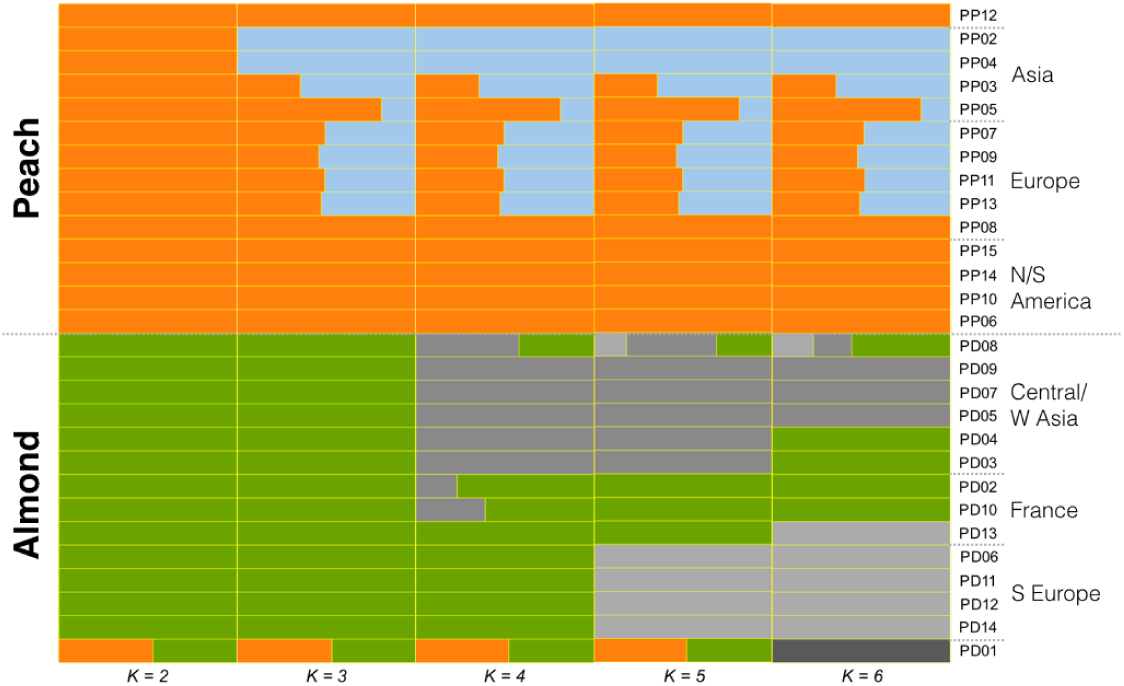
Increasing the assumed clusters to *K* =6 (right) places PD01, the almond-peach F1 hybrid collected from Kharkiv Market, Ukraine, into a unique sub-population. It also shifts the assignments of samples PD13, PD03, and PD04 to different sub-populations, when compared to their assignments in *K* =5 (second from right).

**Figure S5:**
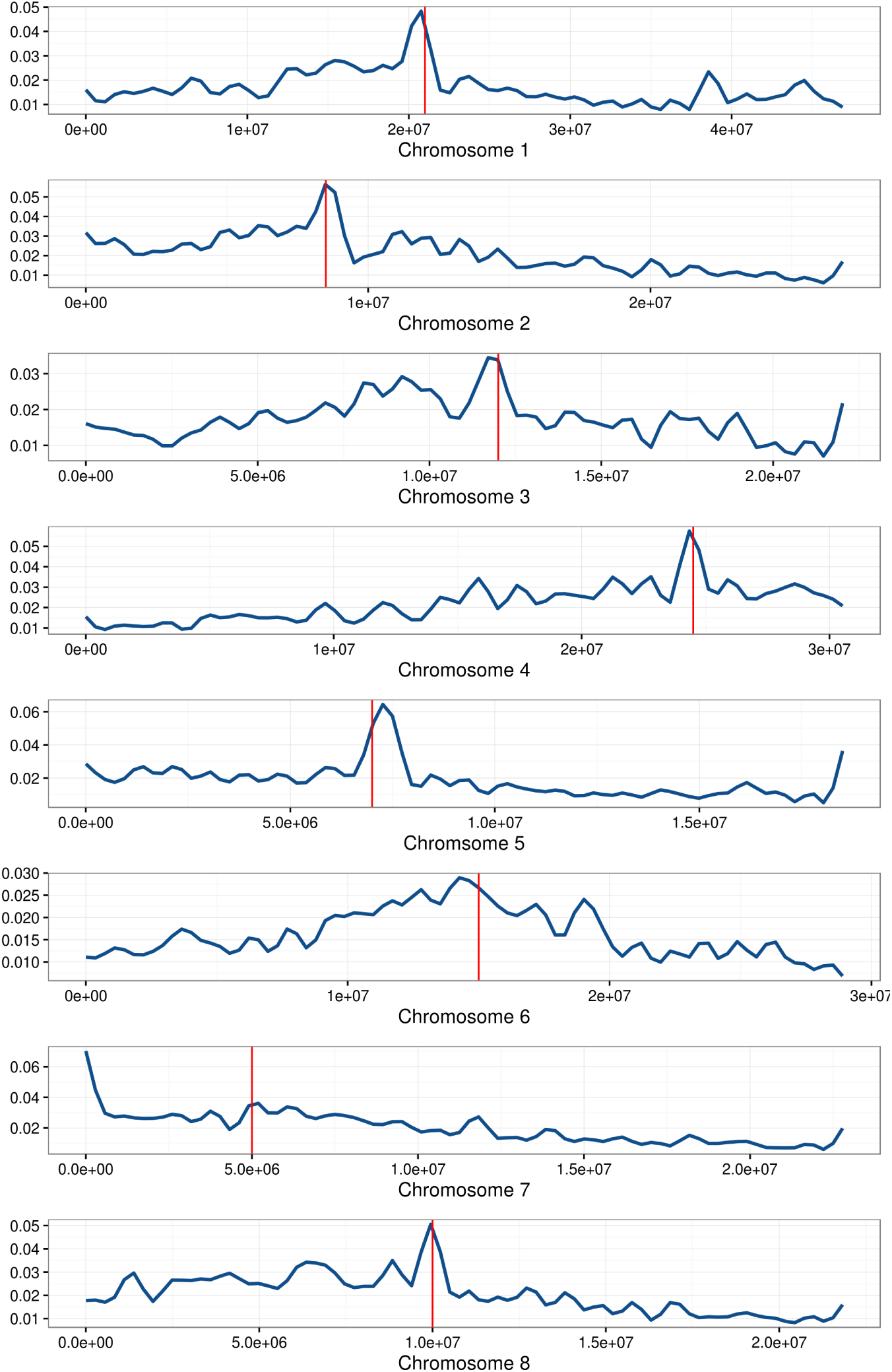
Nucleotide diversity (*θ_π_*) in almond for each chromosome. The vertical red line indicates the approximate location of the centromere.

**Figure S6:**
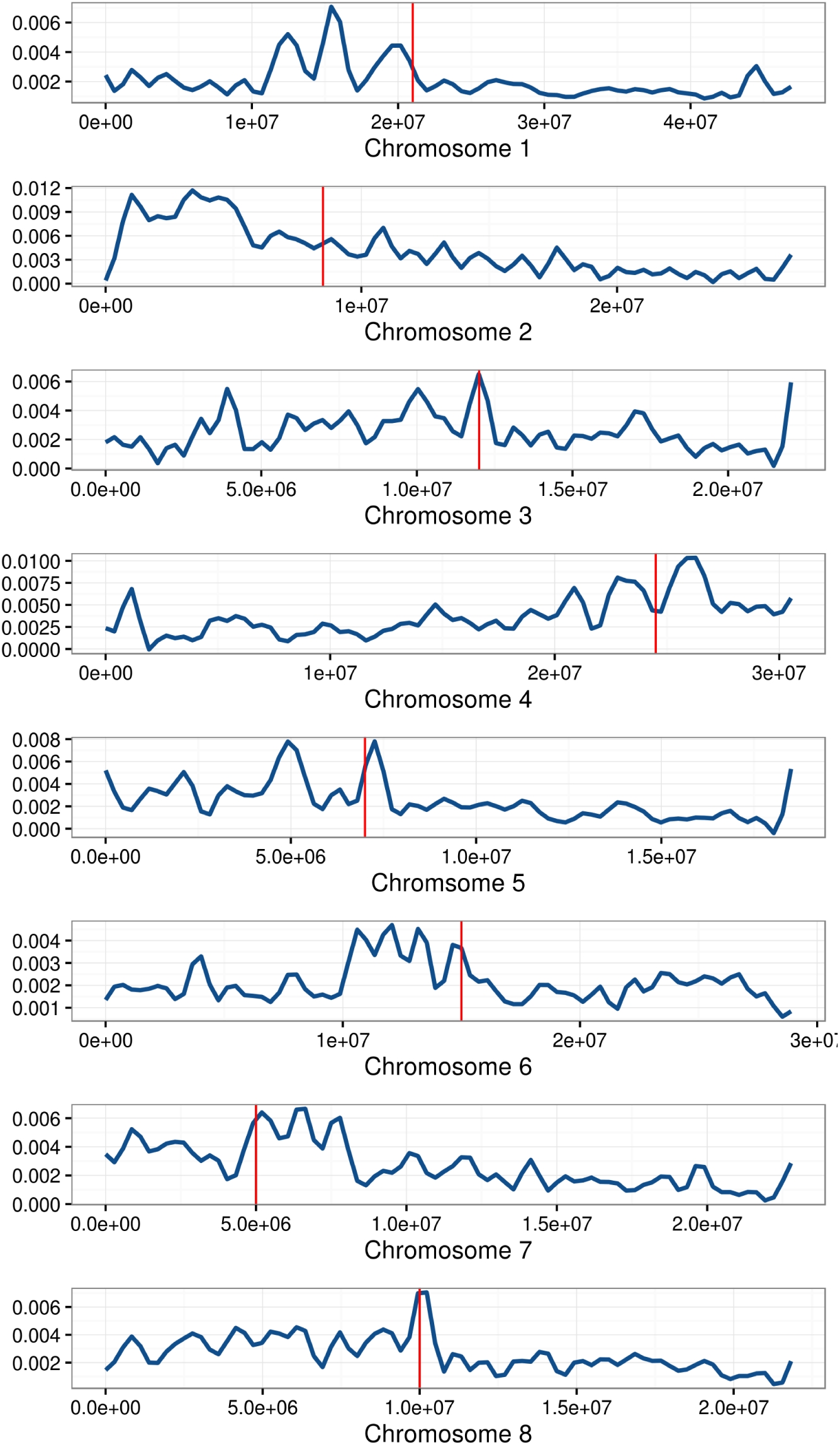
Nucleotide diversity (*θ_π_*) in peach for each chromosome. The vertical red line indicates the approximate location of the centromere.

**Figure S7:**
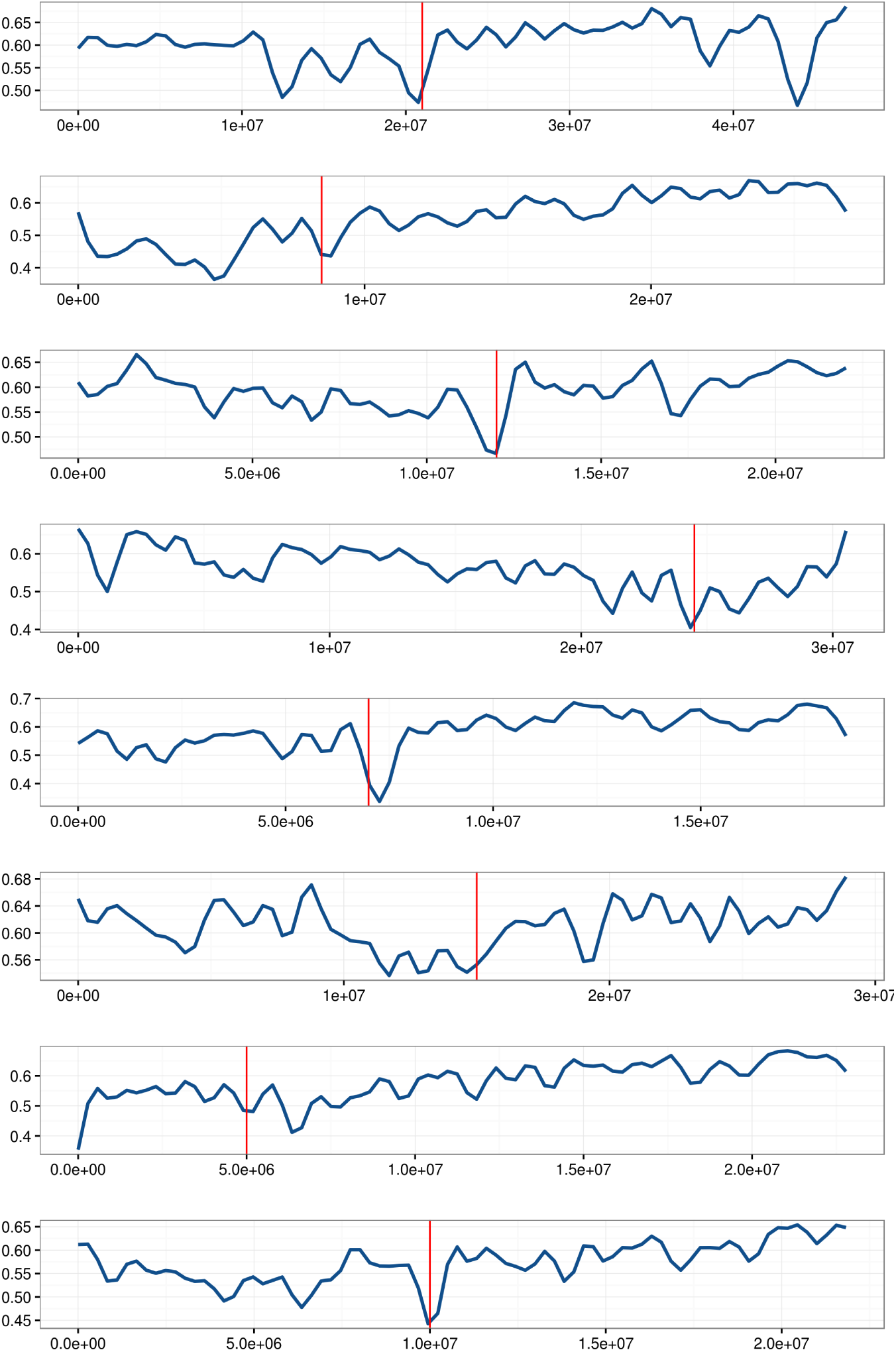
*F*_*ST*_ between almond and peach for each chromosome. The vertical red line indicates the approximate location of the centromere.

**Figure S8:**
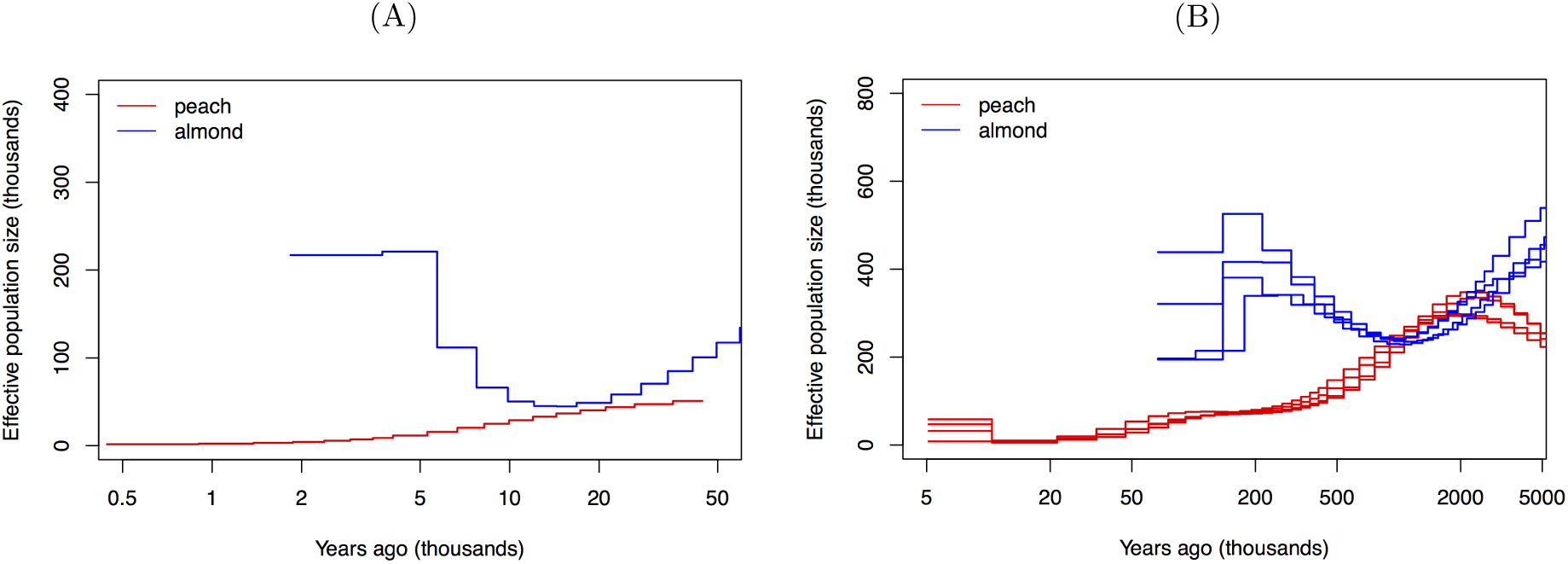
Historical changes in N_*e*_ over time in both almond and peach. Shown are estimates of N_*e*_ for both the (A) recent (*≤* 50 *×* 103 BP) and (B) ancient (*≤* 5000 *×* 103 BP) past.

**Table S1:**
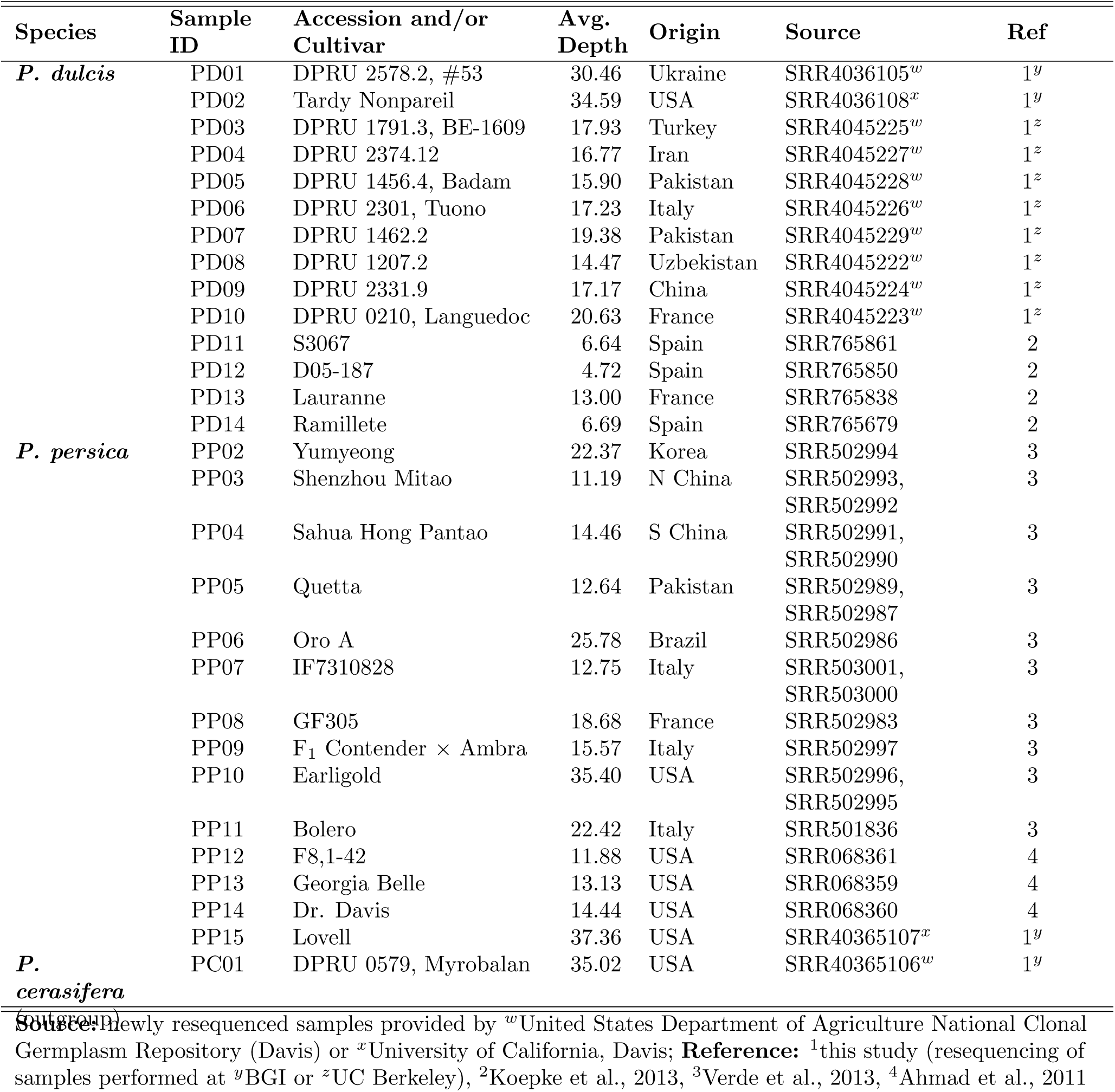
Detailed sample information for *P. dulcis*, *P. persica*, and related species used in analyses.

**Table S2:** RNA-seq data used in expression analyses.

| **SRA Run ID** | **Species** | **Tissue** | **Cultivar** | **Reference** |
| --- | --- | --- | --- | --- |
| SRR2086434 | Peach | fruit mesocarp | Red Pearl | Sanhueza et al. 2015 |
| SRR2086433 | Peach | fruit mesocarp | Red Pearl | Sanhueza et al. 2015 |
| SRR3823906 | Peach | fruit mesocarp | DU-88 | N/A |
| SRR3823907 | Peach | fruit mesocarp | DU-88 | N/A |
| SRR2290949 | Peach | leaf | Jangtaek | Jo et al. 2015 |
| SRR2290951 | Peach | leaf | Mibaek | Jo et al. 2015 |
| SRR1662173 | Peach | leaf | Hongyetao | Wang et al. 2013 |
| SRR1662174 | Peach | leaf | Mantianhong | Wang et al. 2013 |
| SRR2976060 | Almond | ovary | genotype H | Mousavi et al. 2014 |
| SRR2976058 | Almond | anther | genotype H | Mousavi et al. 2014 |

N/A: unable to locate a publication for this data submitted to SRA by the Andres Bello Universidad

**Table S3:** Inbreeding values of peach and almond samples.

| **Peach** | *F* | **Almond** | *F* |
| --- | --- | --- | --- |
| PP02 | 0.072 | PD02 | 0.000 |
| PP03 | 0.222 | PD03 | 0.002 |
| PP04 | 0.116 | PD04 | 0.000 |
| PP05 | 0.001 | PD05 | 0.002 |
| PP06 | 0.533 | PD06 | 0.000 |
| PP07 | 0.081 | PD07 | 0.000 |
| PP08 | 0.737 | PD08 | 0.000 |
| PP09 | 0.000 | PD09 | 0.000 |
| PP10 | 0.064 | PD10 | 0.000 |
| PP11 | 0.000 | PD11 | 0.000 |
| PP13 | 0.000 | PD12 | 0.027 |
| PP14 | 0.176 | PD13 | 0.000 |
| PP15 | 0.557 | PD14 | 0.000 |
| *Mean* | *0.197* | *Mean* | *0.002* |

**Table S4:**
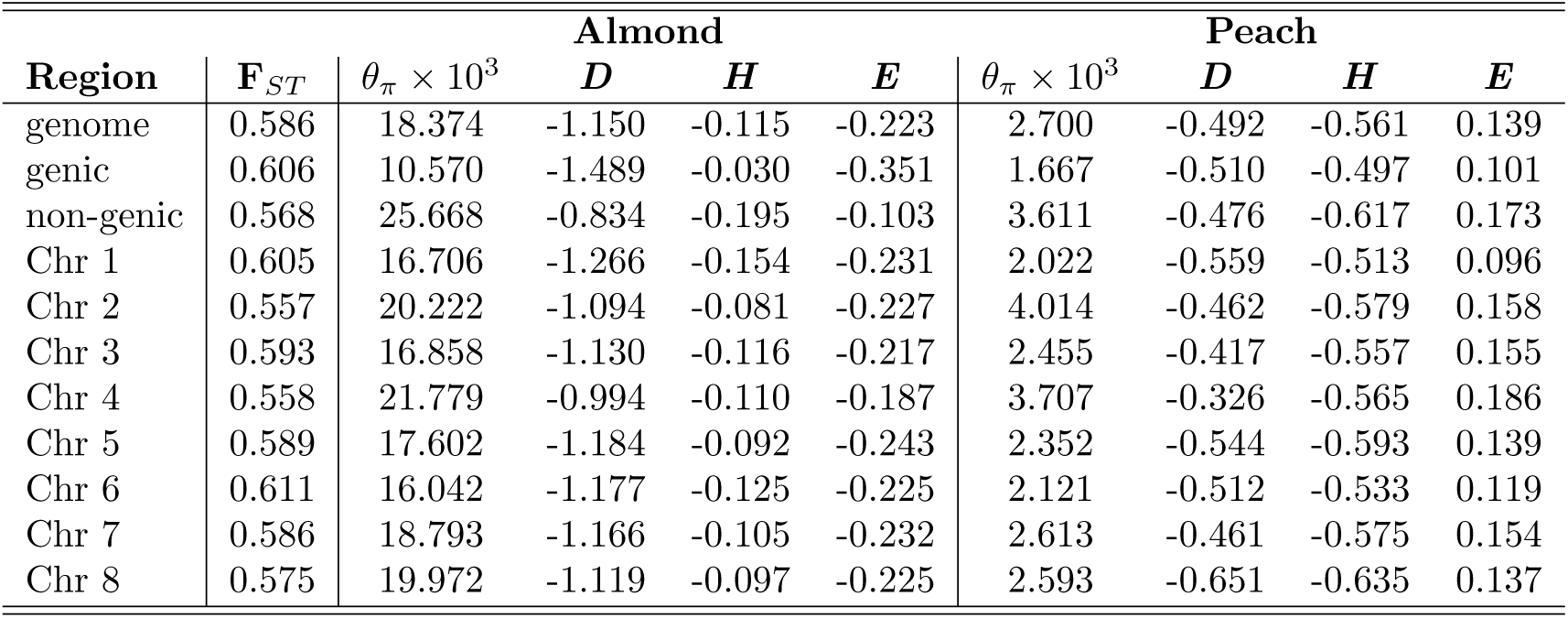
Mean F_*ST*_, diversity statistics, and neutrality test values.

**Table S5:** Significant GO terms for *F*_*ST*_ candidate genes based on top 5% quantile. (Type: F - molecular function; P - biological process)

| GO acc | Type | Term | Query Item | BG Item | p-value | FDR |
| --- | --- | --- | --- | --- | --- | --- |
| GO:0030554 | F | adenyl nucleotide binding | 153 | 2225 | 7.7e-07 | 9.3e-05 |
| GO:0005524 | F | ATP binding | 146 | 2104 | 8.8e-07 | 9.3e-05 |
| GO:0005515 | F | protein binding | 122 | 1634 | 2.5e-07 | 9.3e-05 |
| GO:0001883 | F | purine nucleoside binding | 153 | 2225 | 7.7e-07 | 9.3e-05 |
| GO:0001882 | F | nucleoside binding | 153 | 2226 | 7.9e-07 | 9.3e-05 |
| GO:0032559 | F | adenyl ribonucleotide binding | 146 | 2104 | 8.8e-07 | 9.3e-05 |
| GO:0017076 | F | purine nucleotide binding | 162 | 2437 | 2.3e-06 | 0.00017 |
| GO:0016772 | F | transferase activity, transferring phosphorus-containing groups | 101 | 1347 | 2.4e-06 | 0.00017 |
| GO:0032555 | F | purine ribonucleotide binding | 155 | 2313 | 2.6e-06 | 0.00017 |
| GO:0032553 | F | ribonucleotide binding | 155 | 2313 | 2.6e-06 | 0.00017 |
| GO:0000166 | F | nucleotide binding | 165 | 2531 | 5.4e-06 | 0.00031 |
| GO:0004713 | F | protein tyrosine kinase activity | 62 | 751 | 1.7e-05 | 0.00077 |
| GO:0016798 | F | hydrolase activity, acting on glycosyl bonds | 37 | 363 | 1.6e-05 | 0.00077 |
| GO:0004553 | F | hydrolase activity, hydrolyzing O-glycosyl compounds | 36 | 348 | 1.6e-05 | 0.00077 |
| GO:0004888 | F | transmembrane receptor activity | 26 | 213 | 1.9e-05 | 0.00082 |
| GO:0004872 | F | receptor activity | 26 | 215 | 2.2e-05 | 0.00089 |
| GO:0060089 | F | molecular transducer activity | 31 | 285 | 2.5e-05 | 0.0009 |
| GO:0004871 | F | signal transducer activity | 31 | 285 | 2.5e-05 | 0.0009 |
| GO:0016740 | F | transferase activity | 137 | 2103 | 3.7e-05 | 0.0013 |
| GO:0003964 | F | RNA-directed DNA polymerase activity | 16 | 101 | 5.1e-05 | 0.0016 |
| GO:0034061 | F | DNA polymerase activity | 17 | 115 | 6.5e-05 | 0.002 |
| GO:0016265 | P | death | 38 | 377 | 1.6e-05 | 0.0042 |
| GO:0012501 | P | programmed cell death | 36 | 357 | 2.6e-05 | 0.0042 |
| GO:0023052 | P | signaling | 45 | 486 | 2e-05 | 0.0042 |
| GO:0008219 | P | cell death | 38 | 377 | 1.6e-05 | 0.0042 |
| GO:0006915 | P | apoptosis | 36 | 357 | 2.6e-05 | 0.0042 |
| GO:0006278 | P | RNA-dependent DNA replication | 16 | 101 | 5.1e-05 | 0.0068 |
| GO:0016773 | F | phosphotransferase activity, alcohol group as acceptor | 75 | 1072 | 0.00035 | 0.0097 |
| GO:0003824 | F | catalytic activity | 344 | 6355 | 0.00034 | 0.0097 |
| GO:0016779 | F | nucleotidyltransferase activity | 23 | 217 | 0.00037 | 0.01 |
| GO:0004672 | F | protein kinase activity | 70 | 989 | 0.0004 | 0.01 |
| GO:0006260 | P | DNA replication | 19 | 150 | 0.00016 | 0.01 |
| GO:0002376 | P | immune system process | 20 | 165 | 0.00018 | 0.015 |

| Table S5 – continued from previous page |  |  |  |  |  |  |
| --- | --- | --- | --- | --- | --- | --- |
| GO acc | Type | Term | Query Item | BG Item | p-value | FDR |
| GO:0006955 | P | immune response | 20 | 165 | 0.00018 | 0.015 |
| GO:0045087 | P | innate immune response | 20 | 165 | 0.00018 | 0.015 |
| GO:0016301 | F | kinase activity | 74 | 1089 | 0.00082 | 0.02 |
| GO:0023046 | P | signaling process | 36 | 409 | 0.00032 | 0.021 |
| GO:0023060 | P | signal transmission | 36 | 409 | 0.00032 | 0.021 |
| GO:0005488 | F | binding | 371 | 7025 | 0.0011 | 0.025 |
| GO:0005215 | F | transporter activity | 50 | 682 | 0.0013 | 0.03 |
| GO:0007165 | P | signal transduction | 33 | 378 | 0.00063 | 0.039 |
| GO:0007154 | P | cell communication | 14 | 107 | 0.00083 | 0.048 |

**Table S6:** Number and mean summary statistic values of non-genic and genic windows (NGW and GW, respectively) in the lowest 5% quantile for Tajima’s D, Zeng’s E, Fay & Wu’s H, and *θ_π_* for each species. The same information is shown for windows in the top and bottom 5% quantiles for *F*_*ST*_. Also included are the number of genes represented by genic windows and the ratio of genic to non-genic windows.

| **Statistic** |  | NGW | Mean | GW | Genes | Mean | GW:NGW |
| --- | --- | --- | --- | --- | --- | --- | --- |
| Tajima's D | almond | 17112 | -2.3015 | 203826 | 10365 | -2.3302 | 11.9113 |
|  | peach | 126724 | -2.0946 | 93781 | 6000 | -2.0870 | 0.7400 |
| Zeng's E | almond | 12969 | -0.6501 | 195992 | 11385 | -0.6606 | 15.1123 |
|  | peach | 81258 | -0.5535 | 129763 | 10706 | -0.5494 | 1.5969 |
| Fay & Wu's H | almond | 127429 | -1.0246 | 38095 | 4029 | -1.0325 | 0.2990 |
|  | peach | 107582 | -2.9458 | 105573 | 8526 | -3.0076 | 0.9813 |
| $\theta_\pi$ | almond | 13360 | 0.0033 | 188322 | 9647 | 0.0035 | 14.0960 |
|  | peach | 58287 | 8.6123e-06 | 124818 | 9927 | 7.3075e-06 | 2.1414 |
| $F_{ST}$ | top 5% | 73406 | 0.8716 | 88596 | 7400 | 0.8587 | 1.2069 |
|  | bottom 5% | 51018 | 0.1739 | 35688 | 4692 | 0.1622 | 0.6995 |

**Table S7:** Mann-Whitney U (MWU) and *X*^2^ tests for significance of RNAseq specificity and tissue specific expression of peach fruit, peach leaf, almond ovary, or almond anther and candidate status.

| **Tissue** | **Test** | **$F_{ST}$** | **Almond**<br>**$E$** | **Peach**<br>**$E$** |
| --- | --- | --- | --- | --- |
| **Fruit** (peach) | $\chi^2$ | 0.1284 | <u>0.0052</u> | 0.0530 |
|  | MWU | <u>0.0423</u> | 0.1109 | 0.0987 |
| **Leaf** (peach) | $\chi^2$ | $1.362 \times 10^{-9}$ | $1.52 \times 10^{-15}$ | $6.779 \times 10^{-4}$ |
| | MWU | $3.653 \times 10^{-4}$ | 0.1642 | 0.1216 |
| **Ovary** (almond) | $\chi^2$ | 0.909 | <u>0.0150</u> | <u>0.0074</u> |
|  | MWU | 0.6004 | 0.3680 | 0.6040 |
| **Anther** (almond) | $\chi^2$ | 0.7448 | <u>0.0014</u> | 0.0921 |
|  | MWU | 0.4083 | 0.6753 | 0.5490 |

## References

Ahmad, R., Parfitt, D. E., Fass, J., Ogundiwin, E., Dhingra, A., et al. (2011). Whole genome sequencing of peach (Prunus persica L.) for SNP identification and selection. BMC Genomics, 12(1):569.

Akagi, T., Hanada, T., Yaegaki, H., Gradziel, T. M., and Tao, R. (2016). Genome-wide view of genetic diversity reveals paths of selection and cultivar differentiation in peach domestication. DNA Research, page dsw014.

Alonso, J., Ansón, J., Espiau, M., and Socias i Company, R. (2005). Determination of endodormancy break in almond flower buds by a correlation model using the average temperature of different day intervals and its application to the estimation of chill and heat requirements and blooming date. Journal of the American Society for Horticultural Science, 130(3):308–318.

Aradhya, M. K., Weeks, C., and Simon, C. J. (2004). Molecular characterization of variability and relationships among seven cultivated and selected wild species of *Prunus* L. using amplified fragment length polymorphism. Scientia Horticulturae, 103(1):131–144.

Aranzana, M., Garcia-Mas, J., Carbo, J., and Arús, P. (2002). Development and variability analysis of microsatellite markers in peach. Plant Breeding, 121(1):87–92.

Arús, P., Verde, I., Sosinski, B., Zhebentyayeva, T., and Abbott, A. G. (2012). The peach genome. Tree Genetics & Genomes, 8(3):531–547.

Baird, W. V., Estager, A. S., and Wells, J. K. (1994). Estimating nuclear DNA content in peach and related diploid species using laser flow cytometry and DNA hybridization. Journal of the American Society for Horticultural Science, 119(6):1312–1316.

Bassi, D. and Monet, R. (2008). Botany and taxonomy. In Layne, D. R. and Bassi, D., editors, The Peach: Botany, Production and Uses, chapter 1, pages 1–36. CABI, Oxfordshire, UK.

Beissinger, T. M., Wang, L., Crosby, K., Durvasula, A., Hufford, M. B., et al. (2016). Recent demography drives changes in linked selection across the maize genome. Nature Plants, 2:16084 EP –.

Bray, N. L., Pimentel, H., Melsted, P., and Pachter, L. (2016). Near-optimal probabilistic RNA-seq quantification. Nature Biotechnology, 34(5):525–527.

Browicz, K. and Zohary, D. (1996). The genus Amygdalus L. (Rosaceae): species relationships, distribution and evolution under domestication. Genetic Resources and Crop Evolution, 43(3):229–247.

Byrne, D. (1990). Isozyme variability in four diploid stone fruits compared with other woody perennial plants. Journal of Heredity, 81(1):68–71.

Cao, K., Zheng, Z., Wang, L., Liu, X., Zhu, G., et al. (2014). Comparative population genomics reveals the domestication history of the peach, Prunus persica, and human influences on perennial fruit crops. Genome biology, 15(7):415.

Charlesworth, D. (2003). Effects of inbreeding on the genetic diversity of populations. Philosophical Transactions of the Royal Society of London B: Biological Sciences, 358(1434):1051–1070.

Chin, S.-W., Shaw, J., Haberle, R., Wen, J., and Potter, D. (2014). Diversification of almonds, peaches, plums and cherries–molecular systematics and biogeographic history of Prunus (Rosaceae). Molecular phylogenetics and evolution, 76:34–48.

Delplancke, M., Alvarez, N., Benoit, L., Espíndola, A. I, Joly, H., et al. (2013). Evolutionary history of almond tree domestication in the mediterranean basin. Molecular ecology, 22(4):1092–1104.

Delplancke, M., Alvarez, N., Espíndola, A., Joly, H., Benoit, L., et al. (2012). Gene flow among wild and domesticated almond species: insights from chloroplast and nuclear markers. Evolutionary Applications, 5(4):317–329.

Doebley, J. F., Gaut, B. S., and Smith, B. D. (2006). The molecular genetics of crop domestication. Cell, 127(7):1309–1321.

Doyle, J. J. (1987). A rapid DNA isolation procedure for small quantities of fresh leaf tissue. Phytochem Bull, 19:11–15.

Dozier, W., Powell, A., Caylor, A., McDaniel, N., Carden, E., et al. (1990). Hydrogen cyanamide induces budbreak of peaches and nectarines following inadequate chilling. HortScience, 25(12):1573–1575.

Edwards, S. (1975). The almond industry of Mexico. Master’s thesis, Oregon State University.

Fang, X., Zhang, W., Meng, Q., Gao, J., Wang, X., et al. (2007). High-resolution magnetostratigraphy of the Neogene Huaitoutala section in the eastern Qaidam Basin on the NE Tibetan Plateau, Qinghai Province, China and its implication on tectonic uplift of the NE Tibetan Plateau. Earth and Planetary Science Letters, 258(1):293–306.

Fay, J. C. and Wu, C.-I. (2000). Hitchhiking under positive Darwinian selection. Genetics, 155(3):1405–1413.

Fogle, H. (1977). Self-pollination and its implications in peach improvement. Fruit Varieties Journal.

Fogle, H. W. and Dermen, H. (1969). Genetic and chimeral constitution of three leaf variegations in the peach. Journal of Heredity, 60(6):323–328.

Fresnedo-Ramírez, J., Martínez-García, P. J., Parfitt, D. E., Crisosto, C. H., and Gradziel, T. M. (2013). Heterogeneity in the entire genome for three genotypes of peach [prunus persica (l.) batsch] as distinguished from sequence analysis of genomic variants. BMC genomics, 14(1):750.

Fumagalli, M., Vieira, F. G., Linderoth, T., and Nielsen, R. (2014). ngsTools: methods for population genetics analyses from next-generation sequencing data. Bioinformatics, 30(10):1486–1487.

Gaut, B. S., Díez, C. M., and Morrell, P. L. (2015). Genomics and the contrasting dynamics of annual and perennial domestication. Trends in Genetics.

Glémin, S., Bazin, E., and Charlesworth, D. (2006). Impact of mating systems on patterns of sequence polymorphism in flowering plants. Proceedings of the Royal Society B: Biological Sciences, 273(1604):3011–3019.

Gradziel, T. M. (2011). Origin and dissemination of almond. Horti Rev, 38:23–81.

Gross, B. L., Henk, A. D., Richards, C. M., Fazio, G., and Volk, G. M. (2014). Genetic diversity in Malus× domestica (Rosaceae) through time in response to domestication. American Journal of Botany, 101(10):1770–1779.

Hamrick, J. L., Godt, M. J. W., and Sherman-Broyles, S. L. (1992). Factors influencing levels of genetic diversity in woody plant species. In Population genetics of forest trees, pages 95–124. Springer.

Hanada, T., Watari, A., Kibe, T., Yamane, H., Wϋnsch Blanco, A., et al. (2014). Two novel self-compatible S haplotypes in peach (Prunus persica). Journal of the Japanese Society for Horticultural Science, 83(3):203–213.

Hazzouri, K. M., Escobar, J. S., Ness, R. W., Killian Newman, L., Randle, A. M., et al. (2013). Comparative population genomics in Collinsia sister species reveals evidence for reduced effective population size, relaxed selection, and evolution of biased gene conversion with an ongoing mating system shift. Evolution, 67(5):1263–1278.

Hedrick, U. P., Howe, G. H., Taylor, O. M., and Tubergen, C. B. (1917). The Peaches of New York. JB Lyon Company, Albany, NY.

Holsinger, K. E. and Weir, B. S. (2009). Genetics in geographically structured populations: defining, estimating and interpreting FST. Nature Reviews Genetics, 10(9):639–650.

Jo, Y., Chu, H., Cho, J. K., Choi, H., Lian, S., et al. (2015). De novo transcriptome assembly of two different peach cultivars grown in Korea. Genomics data, 6:260–261.

Kester, D. E. and Sartori, E. (1966). Rooting of cuttings in populations of peach (Prunus persica l.), almond (Prunus amygdalus batsch) and their F1 hybrids. In Proceedings of the American Society for Horticultural Science, volume 88, pages 219–223.

Koepke, T., Schaeffer, S., Harper, A., Dicenta, F., Edwards, M., et al. (2013). Comparative genomics analysis in Prunoideae to identify biologically relevant polymorphisms. Plant Biotechnology Journal, 11(7):883–893.

Korneliussen, T. S., Albrechtsen, A., and Nielsen, R. (2014). ANGSD: analysis of next generation sequencing data. BMC Bioinformatics, 15(1):356.

Ladizinsky, G. (1999). On the origin of almond. Genetic Resources and Crop Evolution, 46(2):143–147.

Li, H. (2013). Aligning sequence reads, clone sequences and assembly contigs with BWA-MEM. arXiv preprint arXiv:1303.3997.

Li, X.-w., Meng, X.-q., Jia, H.-j., Yu, M.-l., Ma, R.-j., et al. (2013). Peach genetic resources: diversity, population structure and linkage disequilibrium. BMC genetics, 14(1):1.

Martínez-Goómez, P., Arulsekar, S., Potter, D., and Gradziel, T. M. (2003). An extended interspecific gene pool available to peach and almond breeding as characterized using simple sequence repeat (SSR) markers. Euphytica, 131(3):313–322.

McKey, D., Elias, M., Pujol, B., and Duputié, A. (2010). The evolutionary ecology of clonally propagated domesticated plants. New Phytologist, 186(2):318–332.

Meyer, R. S., DuVal, A. E., and Jensen, H. R. (2012). Patterns and processes in crop domestication: an historical review and quantitative analysis of 203 global food crops. New Phytologist, 196(1):29–48.

Micheletti, D., Dettori, M. T., Micali, S., Aramini, V., Pacheco, I., et al. (2015). Whole-genome analysis of diversity and SNP-major gene association in peach germplasm. PLOS One, 10(9):e0136803.

Miller, A. J. and Gross, B. L. (2011). From forest to field: perennial fruit crop domestication. American Journal of Botany, 98(9):1389–1414.

Miller, P. J., Parfitt, D. E., and Weinbaum, S. A. (1989). Outcrossing in peach. HortScience, 24(2):359–360.

Molnar, P., Boos, W. R., and Battisti, D. S. (2010). Orographic controls on climate and paleoclimate of Asia: thermal and mechanical roles for the Tibetan Plateau. Annual Review of Earth and Planetary Sciences, 38(1):77.

Mousavi, S., Alisoltani, A., Shiran, B., Fallahi, H., Ebrahimie, E., et al. (2014). De novo transcriptome assembly and comparative analysis of differentially expressed genes in Prunus dulcis mill. in response to freezing stress. PLOS One, 9(8):e104541.

Mowrey, B. D., Werner, D. J., and Byrne, D. H. (1990). Isozyme survey of various species of Prunus in the subgenus Amygdalus. Scientia Horticulturae, 44(3):251–260.

Myles, S., Boyko, A. R., Owens, C. L., Brown, P. J., Grassi, F., et al. (2011). Genetic structure and domestication history of the grape. Proceedings of the National Academy of Sciences, 108(9):3530–3535.

Nei, M. and Li, W.-H. (1979). Mathematical model for studying genetic variation in terms of restriction endonucleases. Proceedings of the National Academy of Sciences, 76(10):5269–5273.

Okie, W. R. (1998). Handbook of peach and nectarine varieties. Performance in the Southeastern United States and index of names. Agriculture Handbook (Washington), (714).

Potter, D. (2011). Prunus. In Wild Crop Relatives: Genomic and Breeding Resources, pages 129–145. Springer.

Rehder, A. (1940). Manual of Cultivated Trees and Shrubs. Macmillan Company, New York.

Ross-Ibarra, J., Morrell, P. L., and Gaut, B. S. (2007). Plant domestication, a unique opportunity to identify the genetic basis of adaptation. Proceedings of the National Academy of Sciences, 104(suppl 1):8641–8648.

Sanhueza, D., Vizoso, P., Balic, I., Campos-Vargas, R., and Meneses, C. (2015). Transcriptomic analysis of fruit stored under cold conditions using controlled atmosphere in Prunus persica cv.”Red Pearl”. Frontiers in plant science, 6.

Schiffels, S. and Durbin, R. (2014). Inferring human population size and separation history from multiple genome sequences. Nature Genetics, 46(8):919–925.

Scorza, R. and Okie, W. R. (1991). Peaches (Prunus). Acta Horticulturae, 290:177–234.

Shen, Z., Ma, R., Cai, Z., Yu, M., and Zhang, Z. (2015). Diversity, population structure, and evolution of local peach cultivars in china identified by simple sequence repeats. Genetics and molecular research: GMR, 14(1):101.

Skotte, L., Korneliussen, T. S., and Albrechtsen, A. (2013). Estimating individual admixture proportions from next generation sequencing data. Genetics, 195(3):693–702.

Slotte, T., Hazzouri, K. M., Ågren, J. A., Koenig, D., Maumus, F., et al. (2013). The *Capsella rubella* genome and the genomic consequences of rapid mating system evolution. Nature Genetics, 45(7):831–835.

Spiegel-Roy, P. (1986). Domestication of fruit trees. Developments in agricultural and managed-forest ecology, 16:201–211.

Su, T., Wilf, P., Huang, Y., Zhang, S., and Zhou, Z. (2015). Peaches preceded humans: Fossil evidence from SW China. Scientific Reports, 5:16794–16794.

Tajima, F. (1989). Statistical method for testing the neutral mutation hypothesis by DNA polymorphism. Genetics, 123(3):585–595.

Tao, R., Watari, A., Hanada, T., Habu, T., Yaegaki, H., et al. (2007). Self-compatible peach (Prunus persica) has mutant versions of the S haplotypes found in self-incompatible Prunus species. Plant Molecular Biology, 63(1):109–123.

Verde, I., Abbott, A. G., Scalabrin, S., Jung, S., Shu, S., et al. (2013). The high-quality draft genome of peach (Prunus persica) identifies unique patterns of genetic diversity, domestication and genome evolution. Nature Genetics, 45(5):487–494.

Vieira, J., Fonseca, N. A., Santos, R. A., Habu, T., Tao, R., et al. (2008). The number, age, sharing and relatedness of S-locus specificities in Prunus. Genetics research, 90(01):17–26.

Wang, J., Street, N. R., Scofield, D. G., and Ingvarsson, P. K. (2016). Natural selection and recombination rate variation shape nucleotide polymorphism across the genomes of three related Populus species. Genetics, 202:1185–1200.

Wang, L., Zhao, S., Gu, C., Zhou, Y., Zhou, H., et al. (2013). Deep RNA-seq uncovers the peach transcriptome landscape. Plant Molecular Biology, 83(4-5):365–377.

Wellington, R., Stout, A. B., Einset, O., and Van Alstyne, L. M. (1929). Pollination of fruit trees. Bulletin of the New York State Agricultural Experiment Station, 577:3–54.

Wright, S. I., Kalisz, S., and Slotte, T. (2013). Evolutionary consequences of self-fertilization in plants. Proceedings of the Royal Society of London B: Biological Sciences, 280(1760):20130133.

Wu, J., Gu, C., Khan, M. A., Wu, J., Gao, Y., et al. (2013). Molecular determinants and mechanisms of gametophytic self-incompatibility in fruit trees of Rosaceae. Critical Reviews in Plant Sciences, 32(1):53–68.

Zeinalabedini, M., Khayam-Nekoui, M., Grigorian, V., Gradziel, T., and Martinez-Gomez, P. (2010). The origin and dissemination of the cultivated almond as determined by nuclear and chloroplast SSR marker analysis. Scientia Horticulturae, 125(4):593–601.

Zeng, K., Fu, Y.-X., Shi, S., and Wu, C.-I. (2006). Statistical tests for detecting positive selection by utilizing high-frequency variants. Genetics, 174(3):1431–1439.

Zheng, Y., Crawford, G. W., and Chen, X. (2014). Archaeological evidence for peach (Prunus persica) cultivation and domestication in China. PLOS One, 9(9):e106595.

Zohary, D., Hopf, M., and Weiss, E. (2012). Domestication of Plants in the Old World. Oxford University Press, Oxford.

